# PSME3 Regulates Migration and Differentiation of Myoblasts

**DOI:** 10.1101/2025.01.08.632048

**Authors:** Kenneth D Kuhn, Ukrae H Cho, Martin Hetzer

**Affiliations:** Institute of Science and Technology Austria; University of North Carolina at Chapel Hill

## Abstract

The acquisition of cellular identity requires large-scale alterations in cellular state. The non-canonical proteasome activator PSME3 is known to regulate diverse cellular processes, but its importance for differentiation remains unclear. Here, we demonstrate that PSME3 binds dynamically to highly active promoters over the course of differentiation. However, loss of PSME3 does not globally affect mRNA transcription. We find instead that PSME3 interacts with the HSP90 co-chaperone NUDC and regulates the levels of several cell adhesion-related proteins, such that loss of PSME3 results in increased cell motility. Further, we show that PSME3 facilitates myoblast differentiation in a proteasome-independent manner. Our findings reveal several new facets of PSME3 functionality and highlight its particular importance for the differentiation of myogenic cells.

## Introduction

Differentiation requires the coordinated restructuring of diverse cellular systems. Critical to this process are changes in nuclear organization, which have long been known to be influenced by the activity of the proteasome (McCann and Tansey, 2014). Indeed, the proteasome has such an intimate role in chromatin dynamics, including the modification of histones and initiation of transcription, that one of its components was originally mistaken for a member of the RNA polymerase II (RNAPII) holoenzyme (Kim et al., 1994; Muratani and Tansey, 2003). The proteasome consists of a proteolytic core capped on either end by a regulatory subunit, the best-studied of which is the ubiquitin-binding 19S complex. However, other subunits have been found to play similarly important roles in regulating nuclear processes. Chief among these is proteasome activator complex subunit 3 (referred to here as PSME3, though also known as PA28γ, 11S REGγ, or Ki nuclear autoantigen) (Cascio, 2021).

PSME3 is a nuclear, homoheptameric protein whose function has been linked to the organization of the nuclear speckle, Cajal body, and promyelocytic leukemia body (Baldin et al., 2008; Cioce et al., 2006; Jonik-Nowak et al., 2018; Zannini et al., 2009). PSME3 has also been shown to interact with several important regulators of the cell cycle including p16, p19, and p21, as well as the transcriptional regulators c-Myc, KLF2, SMURF2, NF-κB, and LATS1/2 (Chen et al., 2007; Li et al., 2015; Nie et al., 2010; Sun et al., 2016; Wang et al., 2018; Xu et al., 2016).

Because of these functions, loss of PSME3 is associated with G1 arrest in cultured cells and impaired growth rates in mice (Barton et al., 2004; Chen et al., 2017; Masson et al., 2001). Conversely, PSME3 is frequently overexpressed in various cancer cell lines and associated with accelerated cell division, metastatic potential, and reduced rates of apoptosis (Lei et al., 2020; Mao et al., 2008). Though PSME3 is often understood to operate through the degradation of target proteins, only a small portion of the PSME3 population is associated with the core proteasome (Fabre et al., 2014; Welk et al., 2016). In fact, several of its functions are carried out in a proteasome-independent manner. Through an unknown mechanism, PSME3 can induce mitotic arrest, regulate p53 levels, and maintain global chromatin compaction even when prevented from interacting with the proteasome (Fesquet et al., 2021; Zannini et al., 2008; Zhang and Zhang, 2008).

While PSME3 performs diverse functions in systems critical for differentiation, the mechanisms by which it drives cell identity acquisition are only beginning to be understood. It was recently found that loss of PSME3 impairs T cell maturation and triggers the differentiation of Th17 cells by altering the cell-surface protein profile of dendritic cells (Barton et al., 2004; Zhou et al., 2020). Additionally, suppression of PSME3 expression biases bone marrow stromal cells towards an adipogenic rather than osteogenic fate, and mice lacking PSME3 display corresponding bone-healing defects (Chen et al., 2024). Given these findings, we hypothesized that PSME3 may be important for the differentiation of other cell types, such as those found in the muscular system.

To investigate this possibility, we used C2C12 myoblasts, which can be differentiated into syncytial myotubes upon withdrawal of serum, to interrogate multiple stages of differentiation (Blau et al., 1983; Yaffe and Saxel, 1977). We find for the first time that PSME3 binds extensively to the chromatin at highly active promoter regions, likely through an association with the protein RNAPII regulator RPRD1A. Surprisingly, however, PSME3 depletion has no global effect on gene expression. Instead, we discover that PSME3 interacts with HSP90 co-chaperone NUDC and regulates the levels of cell adhesion- and migration-related proteins. As a result, myoblasts lacking PSME3 display accelerated migration and impaired myogenesis. Finally, we show that PSME3 regulates this process in a cell-intrinsic, proteasome-independent manner, thereby establishing PSME3 as a novel regulator of myoblast differentiation.

## Results

### PSME3 binds to highly active promoters prior to differentiation

Previous studies have established a role for PSME3 in maintaining the integrity of a variety of nuclear structures. In particular, PSME3 was found to associate with several heterochromatic regions and maintain their compacted state (Fesquet et al., 2021). We sought to extend this work by globally investigating the binding of PSME3 to genomic regions across differentiation. To test whether PSME3 associates with chromatin in C2C12 myoblasts, we performed Cleavage Under Targets & Release Using Nuclease (CUT&RUN) against the endogenous population of PSME3. Surprisingly, we found that PSME3 associates extensively with the chromatin at over 5,000 distinct regions in cycling myoblasts (Fig. 1 A and B, Sup. File 1). A large majority of these peaks were found to be co-positive for the active promoter mark H3K4me3. Furthermore, PSME3 showed an even stronger preference for promoter-proximal regions (<= 1kb) than H3K4me3, mainly at the expense of binding to gene bodies or intergenic regions (Fig. 1 C). Of all transcriptionally active promoters, those bound by PSME3 are on average roughly 50% more active than those that lack PSME3 (Fig. 1 D).

**Figure 1:**
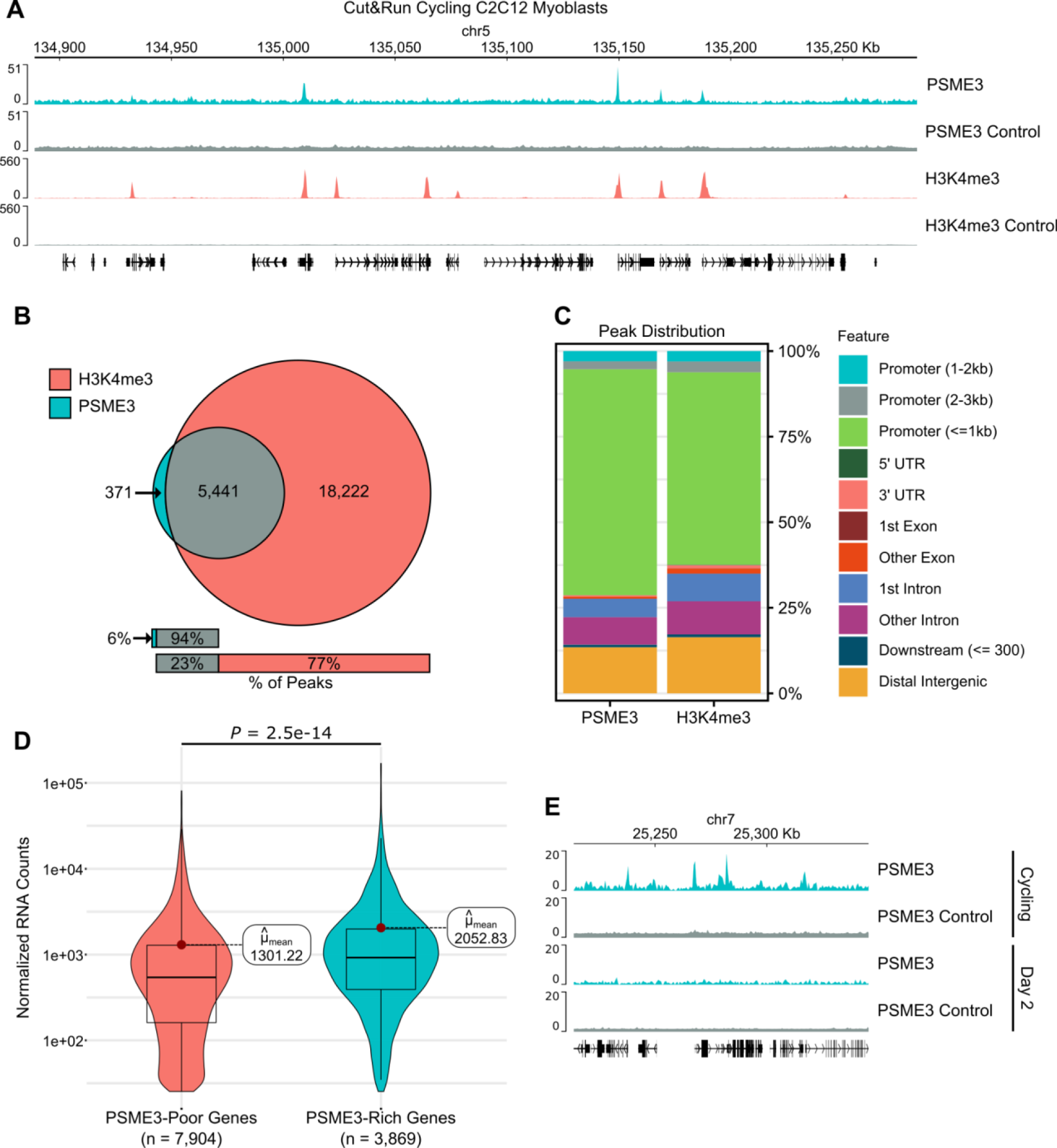
PSME3 dynamically binds to highly active promoters in cycling myoblasts. **A**) CUT&RUN was performed in cycling C2C12 cells using antibodies targeting PSME3 or H3K4me3 to measure their position on the chromatin. Three biological replicates were performed and are represented here together. **B**) Peaks were called using MACS2 for both PSME3 and H3K4me3, and their coincidence was measured. **C**) PSME3 and H3K4me3 CUT&RUN peaks were annotated with ChIPseeker for the type of region in which they reside. **D**) RNA sequencing was performed on cycling C2C12 cells. Active genes that possess a PSME3 peak were separated from those that did not and expression levels were analyzed. Three biological replicates per condition; analyzed with Welch two sample t-test. **E**) CUT&RUN was performed additionally in Day 2 differentiated C2C12 cells using an antibody targeting PSME3. Each track is the result of a single replicate, and the cycling replicate is drawn from those represented in panel (**A**).

To determine whether PSME3 exchanges its binding sites as differentiation progresses, we performed CUT&RUN in myoblasts that had differentiated for two days. In contrast with cycling cells, Day 2 cells displayed a near complete absence of PSME3 at the chromatin (Fig. 1 E). Even when subjected to mild formaldehyde fixation to reduce the lability of transient chromatin interactions, scarcely any increase in the number of PSME3 peaks was observed in cells assayed, analyzed either fresh or after freezing (Fig. S1). In summary, PSME3 binds selectively to highly active promoters in myoblasts, but becomes undetectable by the second day of myogenic differentiation.

### PSME3 interacts with RPRD1A but does not regulate gene expression

To understand what function PSME3 might be serving at the chromatin, we performed co-immunoprecipitation on endogenous PSME3 to identify potential interaction partners (Fig. 2 A). Among the most enriched binding partners were PSME3-interacting protein PSME3IP1, a known PSME3 interactor (Jonik-Nowak et al., 2018), and RPRD1A (Fig. 2 A and B, Sup. File 2). RPRD1A regulates the dephosphorylation of S5 of RNAPII’s C-terminal domain to facilitate progression of the polymerase from the promoter into the gene body (Ali et al., 2019; Ni et al., 2014). Given that S5P is found on polymerases poised to begin transcription and can recruit methyltransferases to deposit H3K4me3 marks on active promoters (Ng et al., 2003), we hypothesized RPRD1A was a potential mediator for PSME3’s chromatin binding activity.

**Figure 2:**
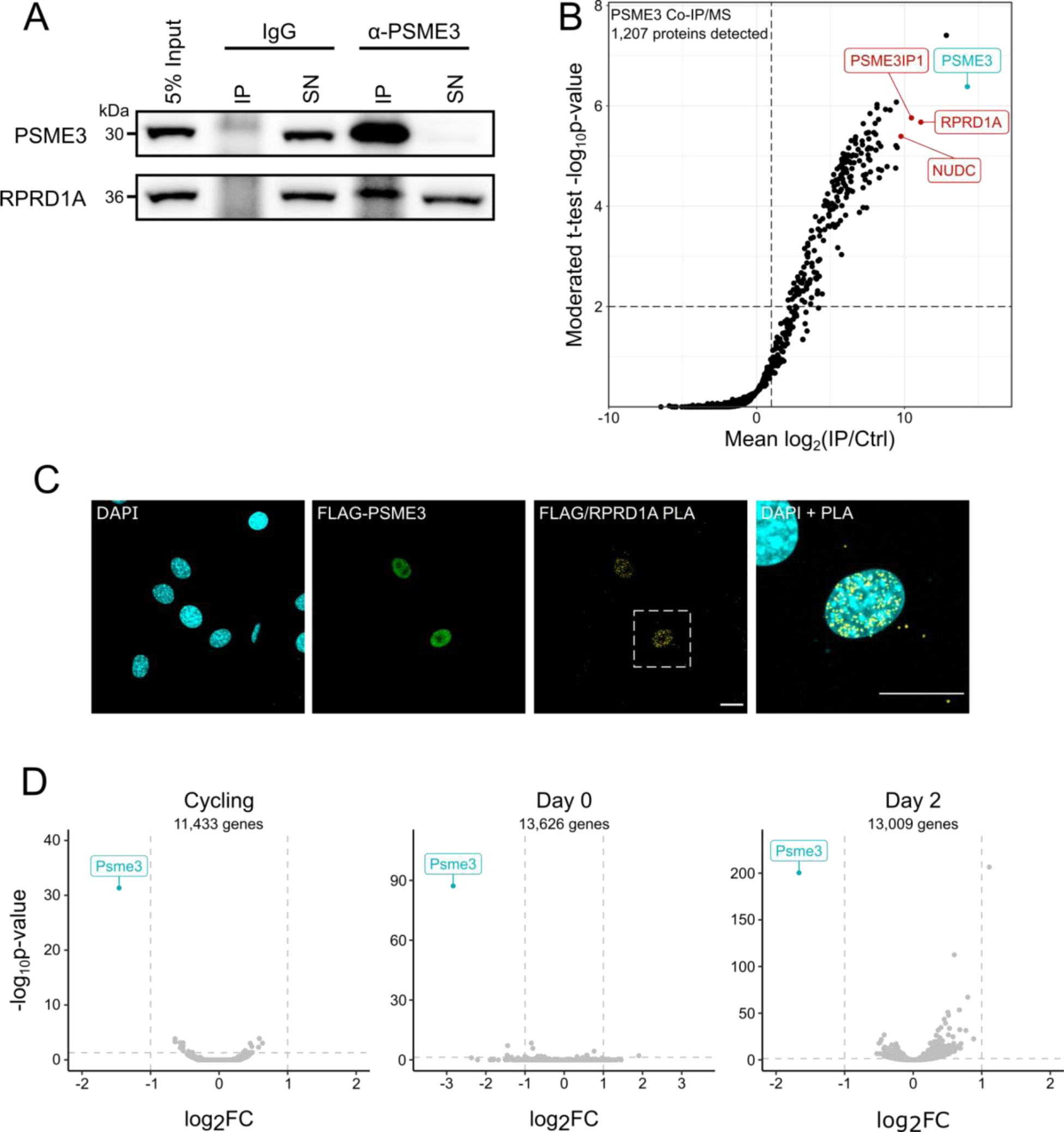
PSME3 interacts with RPRD1A without affecting global gene expression. **A**) PSME3 was precipitated from C2C12 cells while cycling and at days 0 and 1 of differentiation using a PSME3-specific antibody. Efficiency of PSME3 and RPRD1A capture from Day 0 lysates compared to the supernatant (SN) versus that performed with an IgG control was assessed by a Western blot using antibodies targeting PSME3 or RPRD1A. **B**) Label-free quantitative mass spectrometry was performed on the PSME3 co-precipitate across all three timepoints. **C**) Cycling C2C12 cells expressing FLAG-PSME3 were subject to proximity ligation assay (PLA) using primary antibodies targeting either FLAG or RPRD1A. Scale bar is 20 microns in length. **D**) RNA was collected from Cycling, Day 0, or Day 2 C2C12 cells lacking PSME3 and compared to cells treated with scrambled siRNA.

To verify this interaction *in situ*, we utilized a proximity ligation assay (PLA), which produces a fluorescent signal at points where two proteins are removed by a distance of <40 nm. Cycling C2C12 cultures were transfected with a plasmid expressing FLAG-PSME3, and the colocalization of FLAG with RPRD1A was assessed (Fig. 2 C). Cells expressing FLAG-PSME3 showed a high level of PLA signal, while non-transfected cells showed little to no signal. Omission of either antibody similarly abolished the signal (Fig. S2), indicating its specificity.

As alterations in both RPRD1A and PSME3 function have previously been demonstrated to affect gene expression, we asked if PSME3 may be a regulator of transcription during differentiation (Bhatti et al., 2019; Li et al., 2006; Sun et al., 2016; Wang et al., 2018; Wu et al., 2010; Xu et al., 2016). We performed RNA sequencing on PSME3-deficient cells at three stages of differentiation: prior to differentiation (Cycling), fully confluent cells immediately before the withdrawal of serum (Day 0), and cells that have differentiated for two days and begun to form myotubes (Day 2). At each of these time points, we observed no global change in gene expression in cells depleted of PSME3 (Fig. 2 D, Sup. File 3). Taken together, these results suggest that PSME3 interacts with RPRD1A in C2C12 myoblasts, but does not regulate transcription during myogenesis.

### PSME3 regulates cell migration rates and levels of adhesion proteins

During the course of our experiments, we noticed that cells lacking PSME3 were more difficult to detach from the culture plates with trypsin, indicating a greater level of adhesion (data not shown). We therefore turned our attention to the next most enriched protein in the PSME3 immunoprecipitate, NUDC (Fig. 2 B). NUDC was recently identified as a co-chaperone of HSP90 and has been documented to regulate cell migration through the stabilization of several cytoskeletal proteins related to cell adhesion and migration, including cofilin and filamin A (Biebl et al., 2022; Islam et al., 2020; Liu et al., 2021; Zhang et al., 2016).

The observed interaction between NUDC and PSME3 was unexpected, as NUDC is considered to be a predominantly cytoplasmic protein (Islam et al., 2020; Zhang et al., 2016; Zhou et al., 2003; Zhu et al., 2010), while PSME3 is nuclear but may relocalize to the cytoplasm under certain conditions (Carrettiero et al., 2022; Kobayashi et al., 2013; Pecori et al., 2021; Zannini et al., 2008). To better understand the spatial overlap between the two proteins in our system, we first performed live imaging on cycling C2C12 cells that were transfected with constructs expressing PSME3 tagged with either N- or C-terminal GFP, revealing PSME3’s presence in the nucleus (Fig. S3 A). Further, we found that NUDC displayed a strong signal across both cytoplasm and nucleus, where it overlapped with PSME3 staining (Fig. S3 B), indicating that they are well-positioned to interact. To verify this interaction, we performed PLA using antibodies targeting FLAG-PSME3 and NUDC, which specifically revealed the association of the two proteins within the nucleus (Fig. 3 A and S3 C).

**Figure 3:**
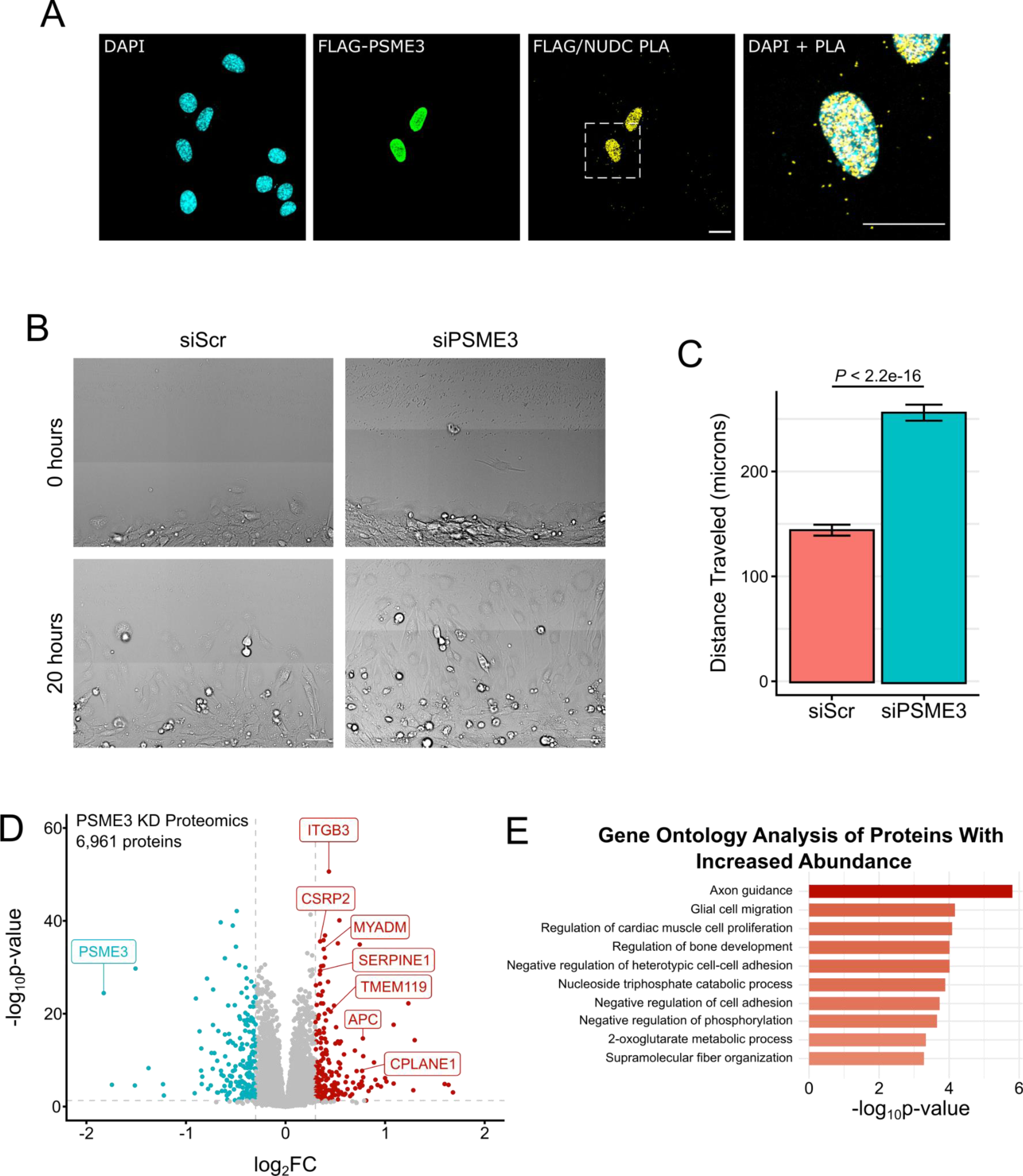
PSME3 associates with NUDC and regulates cell migration. **A**) PLA was performed in cycling C2C12 cells expressing FLAG-PSME3 using antibodies against FLAG and NUDC, and with the respective omission of either or both antibodies as controls. Scale bar is 20 microns in length. **B**) Day 0 confluent C2C12 cells were scratched and allowed to migrate for 20 hours while undergoing continuous brightfield imaging. **C**) Quantification of the distance traveled over 20 hours by individual migrating cells treated with either scrambled (Scr) siRNA or that targeting PSME3. Data was collected with three biological replicates each with two technical replicates of *n* = 15 each and analyzed Welch two sample t-test; error bars show the standard error of the mean. **D**) Day 0 confluent C2C12 cells treated with either scrambled or PSME3-targeting siRNA were subjected to label-free proteomics. The changes in protein abundance are plotted here with decreased-abundance proteins in blue and increased-abundance proteins in red, with several select species involved in cell migration receiving a label. **E**) Gene ontology analysis of increased-abundance proteins; the top ten categories are shown.

To test for a functional interaction between PSME3 and NUDC, we measured the migration rates of fully confluent Day 0 myoblasts in a scratch assay. Cells lacking PSME3 showed a significant increase in migration rate over the control group, consistent with an alteration in cytoskeletal behavior (Fig. 3 B and C). To determine the molecular basis for this change, we performed proteomics on whole-cell lysates from these same cultures. We discovered an increased abundance of several cell adhesion proteins upon PSME3 knockdown, including the Rac1 regulator myeloid-associated differentiation marker (MYADM) and Integrin Beta-3 (ITGB3) (Fig. 3D, Sup. File 4). Gene ontology (GO) analysis of differentially-regulated proteins revealed an increase in species related to cytoskeletal organization, as well as cell migration and adhesion (Fig. 3 E). Taken together, these results indicate that PSME3 interacts with NUDC and regulates both rates of cell migration and levels of cell adhesion proteins.

### Loss of PSME3 impairs myotube formation

We reasoned that if PSME3 affects cell migration, it may also be critical for the differentiation of C2C12 myoblasts, which are particularly sensitive to changes in the cytoskeletal system due to the heavy demands imposed by membrane apposition and cell fusion. To this end, we depleted C2C12 myoblast cells of PSME3 through siRNA treatment prior to serum withdrawal and subsequent differentiation (Fig. 4 A and B). siRNA-treated myoblasts were allowed to differentiate for two days before fixation and staining with antibodies against myogenin, a myogenic transcription factor, and myosin heavy chain (MHC), a major contractile protein found in differentiated myotubes. Widefield imaging revealed a clear deficit in differentiation (Fig. 4 C); cultures lacking PSME3 produced myotubes with fewer nuclei (Fig. 4 D) and possessed a lower fusion index (Fig. 4 E), defined as the percentage of nuclei contained within MHC-positive myotubes.

**Figure 4:**
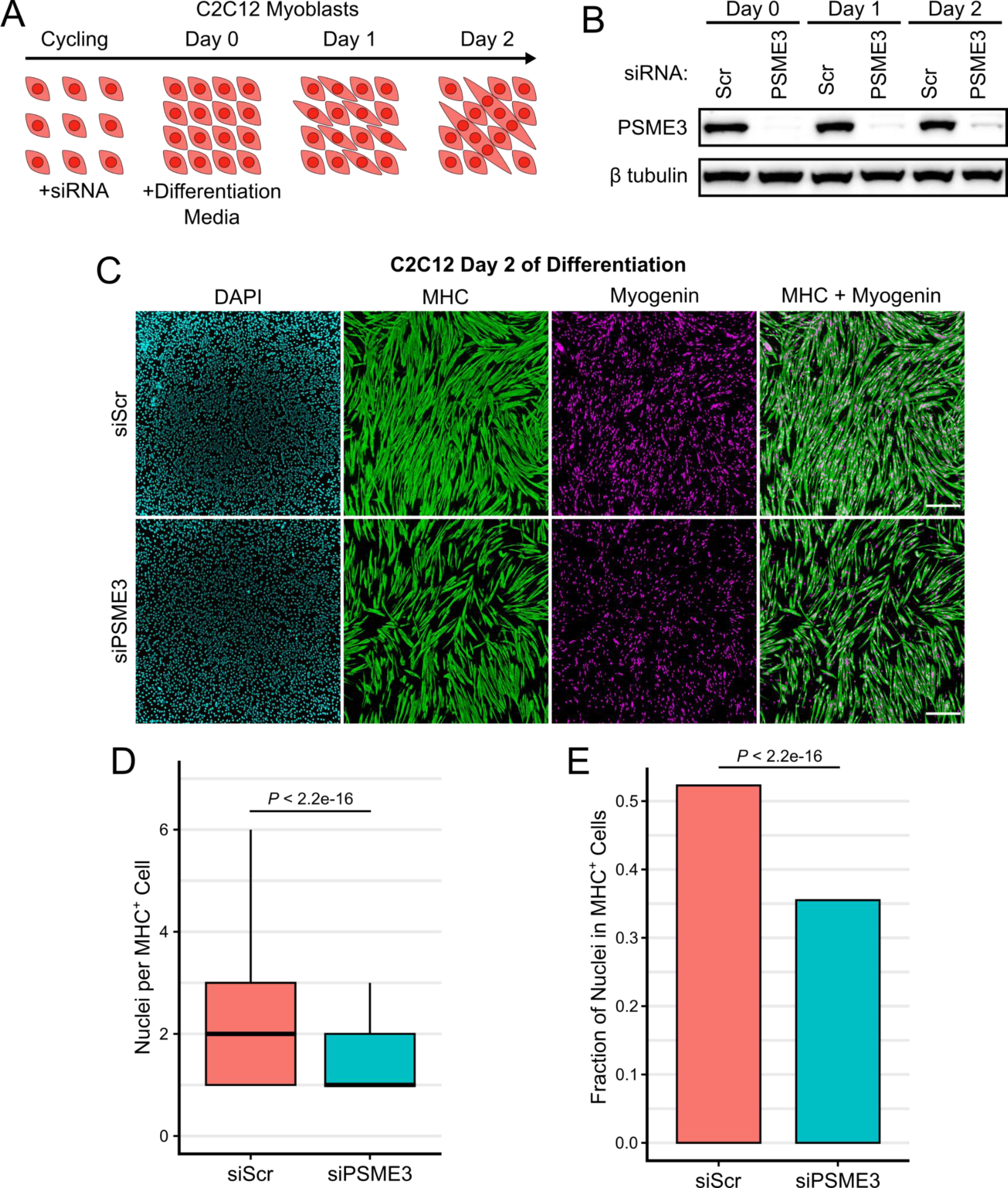
Depletion of PSME3 impairs myotube formation. **A**) Cycling C2C12 cells were treated with siRNA prior to growth to confluency and serum deprivation, which together induce differentiation and myotube formation. **B**) C2C12 cells were treated with scrambled (Scr) or PSME3-targeting siRNA and collected at Day 0, 1, or 2 of differentiation; target protein levels were assessed by Western blot. **C**) C2C12 cells depleted of PSME3 were induced to differentiate for two days and were subjected to immunofluorescence using antibodies targeting MHC and myogenin; scale bar is 200 microns in length. **D**) Immunostained cultures were analyzed for the number of nuclei contained in each MHC-positive cell. Data includes three biological replicates, each comprising two technical replicates of *n* = 250 cells; analyzed by Welch two sample t-test. **E**) Immunostained cultures were analyzed for the percentage of all DAPI-positive nuclei that are contained within MHC-positive cells. Data includes three biological replicates each with two technical replicates with an *n* of roughly 20,000 cells; analyzed by two sample t-test of proportions.

To determine if the effect of PSME3 on myogenesis is dependent on an interaction with the proteasome, we stably expressed FLAG-tagged PSME3 either in its full length or with a deletion of the terminal fourteen amino acids, which renders it unable to associate with the 20S proteasome (Fig. S4 B) (Fesquet et al., 2021; Förster et al., 2005; Ma et al., 1993; Zannini et al., 2008; Zhang and Zhang, 2008). We found that expression of either the full-length or C-terminal deletion mutant was sufficient to rescue the myogenesis phenotype caused by depletion of endogenous PSME3, indicating that PSME3 regulates myoblast differentiation through a proteasome-independent mechanism (Fig. S4 A-C).

Though there were no global alterations in gene expression when PSME3 was depleted, we observed a modest increase in several collagen transcripts on the second day of differentiation (Fig. 2 D). To confirm that the observed differentiation phenotype is the result of a cell-intrinsic effect and not an alteration in the extracellular matrix, we created a mixed culture system of cells treated with scrambled siRNA or that targeting PSME3 and labeled them with a red or blue dye, respectively. If the differentiation phenotype is due to a cell-extrinsic effect, the presence of control cells should improve the differentiation efficiency of those lacking PSME3. However, when the mixed population was induced to differentiate for three days, we observed a selective exclusion of cells lacking PSME3 from mature myotubes (Fig. S4 D and E). These results indicate that PSME3 is necessary for myoblast differentiation in a cell-intrinsic manner.

## Discussion

PSME3 is an elusive and versatile protein possessed of myriad functions, many of which remain to be uncovered. In this study, we reveal several new facets of PSME3’s activity. As the first unbiased study into PSME3’s genome binding properties, we demonstrate a previously unrecognized capacity of PSME3 to associate with RNAPII regulator RPRD1A and highly active promoters. Further, we show a novel interaction of PSME3 with the HSP90 co-chaperone NUDC, through which it may regulate cell migration rates, the levels of cell adhesion-related proteins, and myogenesis.

Though the 19S-containing proteasome has long been known to associate with the chromatin, particularly at highly active gene regions (Auld et al., 2006), the finding that the alternative proteasome cap PSME3 possesses a similar capacity is a novel discovery. We believe that this association may be mediated by an interaction with RPRD1A, which regulates the post-translational modification state of RNAPII and localizes to the promoters of active genes (Liu et al., 2015). This aligns well with our finding that PSME3 mirrors H3K4me3 peaks in topology and localization and is enriched at the most transcriptionally active promoters. This discovery, in combination with a recent study by Fesquet and colleagues showing that PSME3 binds to and is crucial for maintaining heterochromatic regions (Fesquet et al., 2021), suggests that PSME3 possesses broad chromatin-binding activity.

We were surprised, however, to see very little change in gene expression upon PSME3 depletion. PSME3 has previously been demonstrated to influence the expression of individual genes in various cell types (Bhatti et al., 2019; Li et al., 2006; Sun et al., 2016; Wang et al., 2018; Xu et al., 2016), though its loss has no effect on bulk RNA production (Baldin et al., 2008; Cioce et al., 2006). Furthermore, many of the transcriptional regulators documented to be influenced by PSME3, such as p53, MAPK, and YAP1, were unaffected in abundance by the loss of PSME3 in C2C12 myoblasts (Fig. 3 D). Why PSME3 should occupy highly active promoter regions despite having no apparent role in their regulation is unclear at this point. It is possible that PSME3 may be primed to perform some function that is not engaged during the course of differentiation. For example, it is well-established that double strand DNA breaks (DSBs) induce global degradation of RNAPII complexes in a proteasome-dependent manner (Steurer et al., 2022). Further, loss of PSME3 has been found to sensitize cells to radiomimetic treatment and prevent timely repair at DSB sites (Levy-Barda et al., 2011). Thus, it is possible that PSME3 plays a role in degrading RNAPII in the case of DNA damage. Alternatively, PSME3 may respond to other types of cellular insults, such as it has been shown to do in the context of hyperosmotic stress (Carrettiero et al., 2022), and under other such circumstances may be capable of adopting a gene regulatory role.

In addition to DNA binding, we observed a role for PSME3 in maintaining the levels of cell migration-related proteins, as well as an interaction with the co-chaperone NUDC, which mediates cargo transfer between HSP70 and HSP90 (Biebl et al., 2022). NUDC was originally identified in the filamentous fungus *Aspergillus nudilans* for its role in regulating the microtubule-based migration of nuclei within the cell (Fu et al., 2016; Osmani et al., 1990). Additionally, a role for NUDC in regulating cell migration has been demonstrated in diverse cell types through stabilization of several migration-related proteins (Aumais et al., 2001; Cappello et al., 2011; Islam et al., 2020; Liu et al., 2021; Zhang et al., 2016). Indeed, we observed that depletion of PSME3 in C2C12 cells resulted in an increase in the rate of cell migration. Because an improper balance of migration regulators is sufficient to disrupt myogenesis, we suspect that these alterations in cell migration are functionally related to the reduced efficiency of differentiation that we observe in our system (Lehka and Rędowicz, 2020). While we establish PSME3’s independence of the proteasome for differentiation, how PSME3 might interact with NUDC to achieve these functions remains unclear. One possibility is that NUDC aids in the handoff of protein cargo from the chaperone system to the proteasome, a role that has been demonstrated for several members of the Bcl2-associated athanogene (BAG) family of proteins, including BAG2, which is known to associate with both HSP70 and PSME3 under stress conditions (Abildgaard et al., 2020; Carrettiero et al., 2022).

In summary, our results establish PSME3 as an important regulator of myogenesis and will facilitate future research into the contribution of this protein to the acquisition of cell identity.

## Materials and methods

### Plasmids

Constructs were created by insertion of PSME3 into pcDNA3.1(+)-C-eGFP or pcDNA3.1(+)-N-eGFP for live imaging, and pcDNA3.1(+)-N-DYK or pcDNA3.1(+)-C-DYK for FLAG-tagged experiments. PSME3 sequence (accession NM_011192.4) was obtained from GenScript.

### Antibodies

The following antibodies were used: PSME3 (rabbit polyclonal, BML-PW8190, Enzo Life Sciences, 1:6,000 IF, 1:200 CUT&RUN) (rabbit polyclonal, 38-3800, Thermo Fisher Scientific, 1:125 WB, 1:25 IP); β-Tubulin (rabbit monoclonal 9F3, #2128, Cell Signaling Technology; 1:1,000 Western blot); Myosin 4 (mouse monoclonal MF20, 14-6503-82, Thermo Fisher Scientific; 1:100 IF); Myogenin (mouse monoclonal F5D, 14-5643-82, Thermo Fisher Scientific, 1:250 IF); FLAG (mouse monoclonal M2, F1804, Sigma-Aldrich, 1:500 IF and PLA, 1:1,000 Western Blot); RPRD1A (rabbit polyclonal, HPA040602, Sigma-Aldrich, 1:50 PLA, 1:250 Western Blot); NUDC (rabbit polyclonal, 10681-1-AP, Proteintech, 1:25 IF and PLA); H3K4me3 (rabbit monoclonal C42D8, #9751, Cell Signaling Technology, 1:50 CUT&RUN).

### Cell culture and transfection

C2C12 cells obtained from the American Type Culture Collection (ATCC) were cultured in DMEM with 20% FBS. Cells were allowed two passages (four days) after thawing before being subjected to any experiments. No cells over passage nine were used for any experiment.

For transfection with siRNA, cycling C2C12 cells were reverse transfected on two consecutive days with Lipofectamine RNAiMAX transfection reagent (Thermo Fisher Scientific) and either SMARTpool siRNA from Horizon Discovery targeting PSME3 (L-062727-01-0005) or a non-targeting control pool (DH-D-001810-10-20) in OptiMEM. Cells were allowed to recover for two days before being collected or induced to differentiate. For differentiation into myotubes, cells were grown to full confluency, washed once with PBS, and switched to DMEM with 2% horse serum, designated as Day 0. DIfferentiation media was refreshed every 48 hours.

For transfection with plasmids, cells were grown to 80% confluency and were treated with a mixture of Lipofectamine 2000 (Thermo Fisher Scientific, 11668019) and the plasmid of interest. Cells were passaged on the following day and examined on the day thereafter for expression.

### Immunofluorescence and proximity ligation assay

Two methods were used for immunofluorescence. In the first method, cells were fixed with 4% formaldehyde in PBS for five minutes at room temperature. They were then incubated with a blocking/permeabilization buffer containing 0.1% Triton X-100, 0.02% SDS, and 10 mg/mL BSA in PBS. The cells were incubated in primary antibody in the blocking solution for ninety minutes, followed by three washes in PBS, an incubation with fluorescent secondary antibodies in the blocking solution, then an additional three washes in PBS before mounting in Everbrite Mounting Medium with DAPI (Biotium, 23002). This method was used for all imaging experiments that did not involve staining for NUDC.

In the second method, fixation was performed by incubation with 4% formaldehyde in PBS for five minutes at room temperature, followed by eight minutes in cold 100% methanol at −20℃. The blocking/permeabilization buffer was composed of 0.1% saponin with 10 mg/mL BSA, which was also used as a diluent for the primary and secondary antibody. This method is otherwise identical to the first method, and was used exclusively for experiments requiring staining of NUDC.

The proximity ligation assay was performed with the DuoLink In Situ Orange Starter Kit (Sigma-Aldrich, DUO92102-KT) according to the manufacturer’s instructions with the following exceptions: permeabilization/blocking and primary antibody dilution/incubation was performed with the buffers and according to the principles described in the immediately preceding paragraphs. Negative controls included several wells in which one or both of the primary antibodies were omitted, while all other components remained.

Imaging of mature myotubes as well as cell migration was performed with a Nikon Ti2-E Widefield Fluorescence Microscope using a 20x objective, while all other imaging experiments utilized a Leica SP8 Laser Confocal Microscope using a 63x oil immersion objective. All imaging experiments were performed in µ-Slide 8 Well ibiTreat IBIDI chambers.

### Retrovirus production

The coding sequence of PSME3 was FLAG-tagged and cloned into the pQCXIB vector, with which 80% confluent HEK293T cells were transfected along with the pCL-Ampho retrovirus packaging vector in equal proportions. The media was changed at 24 hours after transfection and virus-containing conditioned media was collected an additional 24 hours thereafter. After being passed through a 0.45 micron filter, the conditioned media was combined 1:1 with fresh growth media and incubated with C2C12 cells for 24 hours before exchange with fresh growth media. At 48 hours after infection, cells were selected with 1-2 ug/mL of puromycin until cell death ceased and control cells had uniformly perished. For knockdown experiments, siRNA targeting the 3’-UTR was used to deplete the endogenous copy of PSME3 while sparing the tagged variants.

### Cell mixing

Cells were treated with siRNA as described above, and on the day prior to differentiation were stained with either CellTracker Deep Red Dye (1mM solution at 1:250; Thermo Fisher Scientific, C34565) or CellTrace Violet Dye (5mM at 1:500; Thermo Fisher Scientific, C34571) according to the manufacturer’s instructions and as previously described (Zhang et al., 2017). After staining, cells were counted and either plated in isolation, or in a 1:1 mixture. Differentiation was induced on the following day, and incorporation into myotubes was assessed on day 3 after fixing and staining for MHC. Due to the level of cell death normally observed during differentiation, blue nuclei indicating cells lacking PSME3 were counted by hand. The fraction contained in MHC-positive structures was quantified and compared between the separate conditions.

### Image analysis

The number of myogenin-positive nuclei within each MHC-positive structure was performed manually in ImageJ (Schindelin et al., 2012). The total fraction of nuclei contained within MHC-positive structures was performed by thresholding the MHC channel to create a binary image, followed by watershedding to fill in the gaps within the cell. DAPI-positive nuclei were similarly converted to a binary image, and the MHC image was used as a mask to subtract nuclei not contained within. The total number of nuclei remaining were quantified using the Analyze Particles function of ImageJ.

Cell migration was measured manually by displacement of the center of the nucleus over two hour intervals. The sum of these displacements over ten such intervals (20 hours total) was summed for each individual cell. Cells were selected at hour zero only if they were clearly visible, and were discarded if, during the course of the measurement, they divided or died.

### Immunoblotting

Cells were collected by trypsinization and counted. 200 uL of RIPA buffer (50mM Tris-HCl pH 7.5, 150mM NaCl, 1% Triton X-100, 0.5% sodium deoxycholate, 0.1% SDS, and 1x Pierce Protease Inhibitor Tablet (Thermo Fisher Scientific, A32963)) was added per 1e6 cells, which were incubated for 30 minutes at 4℃ with rotation. Lysates were then briefly sonicated (three five second pulses at approximately 1 Watt) before being clarified by centrifugation for 8 minutes at 10,000 RPM at 4℃ on a tabletop centrifuge. 4x Sample Loading Buffer (Biorad, 1610747) with beta mercaptoethanol was added to the lysates, which were then incubated at 95℃ for five minutes.

Samples were loaded into 4-12% gradient gels (Bolt™ Bis-Tris Plus Mini Protein Gels, Thermo Fisher Scientific, NW04122BOX) and run in Bolt MOPS SDS (Thermo Fisher Scientific, B000102). Proteins were transferred using the Biorad Trans-Blot Turbo Transfer System and were blocked in TBS-T plus 5% milk for one hour before being incubated overnight with primary antibodies in the same solution. Membranes were washed three times in TBS-T before probing with secondary antibodies conjugated with HRP targeting either mouse (Thermo Fisher Scientific, 31430) or rabbit (Thermo Fisher, G-21234). Membranes were incubated with SuperSignal West Pico PLUS Chemiluminescent Substrate (Thermo Fisher Scientific, 34580) before imaging.

### Immunoprecipitation

Cells were collected by trypsinization, washed once with PBS, and lysed for 30 minutes at 4℃ with rotation in an immunoprecipitation (IP) buffer containing 20 mM Tris-HCl pH 7.5, 100mM NaCl, 1% NP40, 1mM EDTA, and 1x Pierce Protease Inhibitor Tablet (Thermo Fisher Scientific, A32963). 200 uL of IP buffer was used per 1e6 cells, and 2.5e6 cells were used for each immunoprecipitation. Cells were clarified by centrifugation for 8 minutes at 10,000 RPM at 4℃ on a tabletop centrifuge. Primary antibodies were added directly into the clarified supernatant at 2.8 ug per 1e6 cells, with an isotype IgG (Biotechne, AB-105-C) used as a control. This solution was incubated overnight at 4℃ with rotation. The following morning, 50 uL of Protein A Dynabeads (Thermo Fisher Scientific, 10008D) was added directly to the solution, which was then incubated for two hours at 4℃ with rotation. Beads were removed from solution with a magnet and washed three times by resuspension with 200 uL of IP buffer. Beads were then collected in 100 uL of IP buffer and transferred to a clean tube before being resuspended in 3x Sample Loading Buffer (Biorad, 1610747) with beta mercaptoethanol and incubated for five minutes at 95℃. The supernatant was collected and frozen on dry ice before being stored at −80℃. Efficacy of the precipitation was assessed via immunoblot as described above, but using a protein A-HRP conjugate (Millipore, 18-160) instead of the usual secondary antibodies to avoid detection of eluted IgG.

### CUT&RUN

CUT&RUN was performed using the kit provided by Cell Signaling Technology (#86652) according to the manufacturer’s instructions, but with the following modifications. Three hundred thousand cells were used per condition. Further, incubation was performed not overnight at 4℃ as the manufacturer recommends, but rather for 30 minutes at room temperature to limit the level of cell death. Cells, unfixed, were frozen prior to assaying in 10% DMSO in FBS unless otherwise indicated.

Libraries were prepared using the NEBNext Ultra™ II DNA Library Prep Kit for Illumina (E7645L) with primers from the NEBNext Multiplex Oligos for Illumina (Dual Index Primers Set 1, E7600S) according to the manufacturer’s instructions. Prepared libraries were sent to Novogene for sequencing. Raw reads were cleaned with Trim Galore and aligned with Bowtie2, whereafter peaks were called with MACS2 (Babraham, 2016; Langmead and Salzberg, 2012; Zhang et al., 2008).

### RNA Sequencing

Cells were collected directly from the plate by washing once with PBS followed by addition of Trizol reagent (Thermo Fisher Scientific, 15596026). The samples were incubated in the plate for three minutes at room temperature prior to freezing in dry ice and long term storage at −80℃. RNA was isolated from the sample using a chloroform extraction followed by purification with the RNeasy kit from Qiagen. Samples were submitted to Novogene for library preparation and sequencing. RNA reads were aligned with RNA STAR and comparative gene expression analysis was performed with DESeq2 (Dobin et al., 2013; Love et al., 2014).

### Mass Spectrometry Sample preparation

Samples in 3x Laemmli buffer were first cleaned up by SP3 using a commercial kit (PreOmics GmbH, 50 mg of beads per sample), then processed using the iST kit (PreOmics GmbH) according to the manufacturer’s instructions. Tryptic digestion was stopped after 1 h and cleaned-up samples were vacuum dried. Finally, samples were re-dissolved by 10 min sonication in the iST kit’s LC LOAD buffer.

### LC-MS/MS analysis

Samples were analyzed by LC-MS/MS on a nanoElute 2 nano-HPLC (Bruker Daltonics) coupled with a timsTOF HT (Bruker Daltonics), concentrated over a “Thermo Trap Cartridge 5mm”, then bound to a PepSep XTREME column (1.5 µm C18-coated particles, 25 cm * 150 µm ID, Bruker P/N 1893476) heated at 50°C and eluted over the following 90 min gradient: solvent A, MS-grade H_₂_O + 0.1% formic acid; solvent B, 100% acetonitrile + 0.1% formic acid; constant 0.60 nL/min flow; B percentage: 0 min, 2%; 90 min, 30%, followed immediately by a 8 min plateau at 95%. MS method: M/Z range = 99.993933-1700 Th, ion mobility range = 0.6-1.6 1/K0; transfer time = 60 µs, pre-pulse storage time = 12 µs, enable high sensitivity modus = off, ion polarity = Positive, scan mode = MS/MS (Pasef); TIMS parameters: ramp time = 100 ms, accumulation time = 100 ms; PASEF parameters: ms/ms scans = 10, total cycle time = 1.167166 s, charge range = 0-5, intensity threshold for scheduling = 2000, scheduling target intensity = 15000, exclusion release time = 0.4 min, reconsider precursor switch = on, current/previous intensity ratio = 4, exclusion window mass width = 0.015 m/z, exclusion window v·s/cm^²^ width = 0.015 V*s/cm^²^.

### LC-MS/MS Data analysis

Raw files were searched in FragPipe version 20.0 against a *Mus musculus* proteome sourced from UniprotKB. Fixed cysteine modification was set to +57.02146 (Cysteine). Variable modifications were set to +15.9949 (Methionine), +42.0106 (protein N-term), +79.96633 (STY) and −17.0265 (Gln -> pyroGlu). Peptide identifications were validated using Percolator. Results were filtered in Philosopher at protein level at FDR 1%. MS1-level peptide quantitation was performed using IonQuant with match-between-runs turned on.

FragPipe’s output was re-processed using in-house R scripts, starting from the psm.tsv tables. MS1 intensities were re-normalized to the median. The long format psm.tsv table was consolidated into a wide format peptidoforms table, summing up quantitative values where necessary. Missing values were imputed using two different strategies: i) the KNN (K-Nearest Neighbours) method for Missing-At-Random values within sample groups, and ii) the QRICL (Quantile Regression Imputation of Left-Censored data) method for Missing-Not-At-Random values. Peptidoform intensity values were re-normalized as follows: 1 and 2. Peptidoform-level ratios were then calculated. Protein groups were inferred from observed peptides, and quantified using an in-house algorithm which: i) computes a mean protein-level profile across samples using individual, normalized peptidoform profiles (“relative quantitation” step), ii) following the best-flyer hypothesis, normalizes this profile to the mean intensity level of the most intense peptidoform (“unscaled absolute quantitation” step); for protein groups with at least 3 unique peptidoforms, only unique ones were used, otherwise razor peptidoforms were also included; Phosphopeptidoforms and their unmodified counterparts were excluded from the calculations. Estimated expression values were log10-converted and re-normalized using the Levenberg-Marquardt procedure. Average log10 expression values were tested for significance using a one-sided moderated t-test per samples group (limma). Significance thresholds were calculated using the Benjamini-Hochberg procedure for False Discovery Rate values of 10%, 20% and 30%. FRegulated protein groups were defined as those with a significant P-value and a log2 ratio greater than 1 for immunoprecipitation experiments, and a log2 ratio greater than 0.3 for proteomics experiments. GO terms enrichment analysis was performed using Metascape Gene Annotation & Analysis Resource (Zhou et al., 2019).

## Acknowledgements

The authors would like to thank Dr. Armel Nicholas, Dr. Ewelina Dutkiewicz, and the remaining staff of the ISTA Mass Spectrometry Facility for their assistance, as well as Dr. Lorenzo Puri and the members of his lab for invaluable discussions relating to the project, and Saki for their clarity of thought and insight. This research was further supported by the Lab Support Facility and the Imaging and Optics Facility of ISTA.

## Conflict of Interests

The authors declare that they have no conflict of interest.

---

**Supplementary Figure 1:**
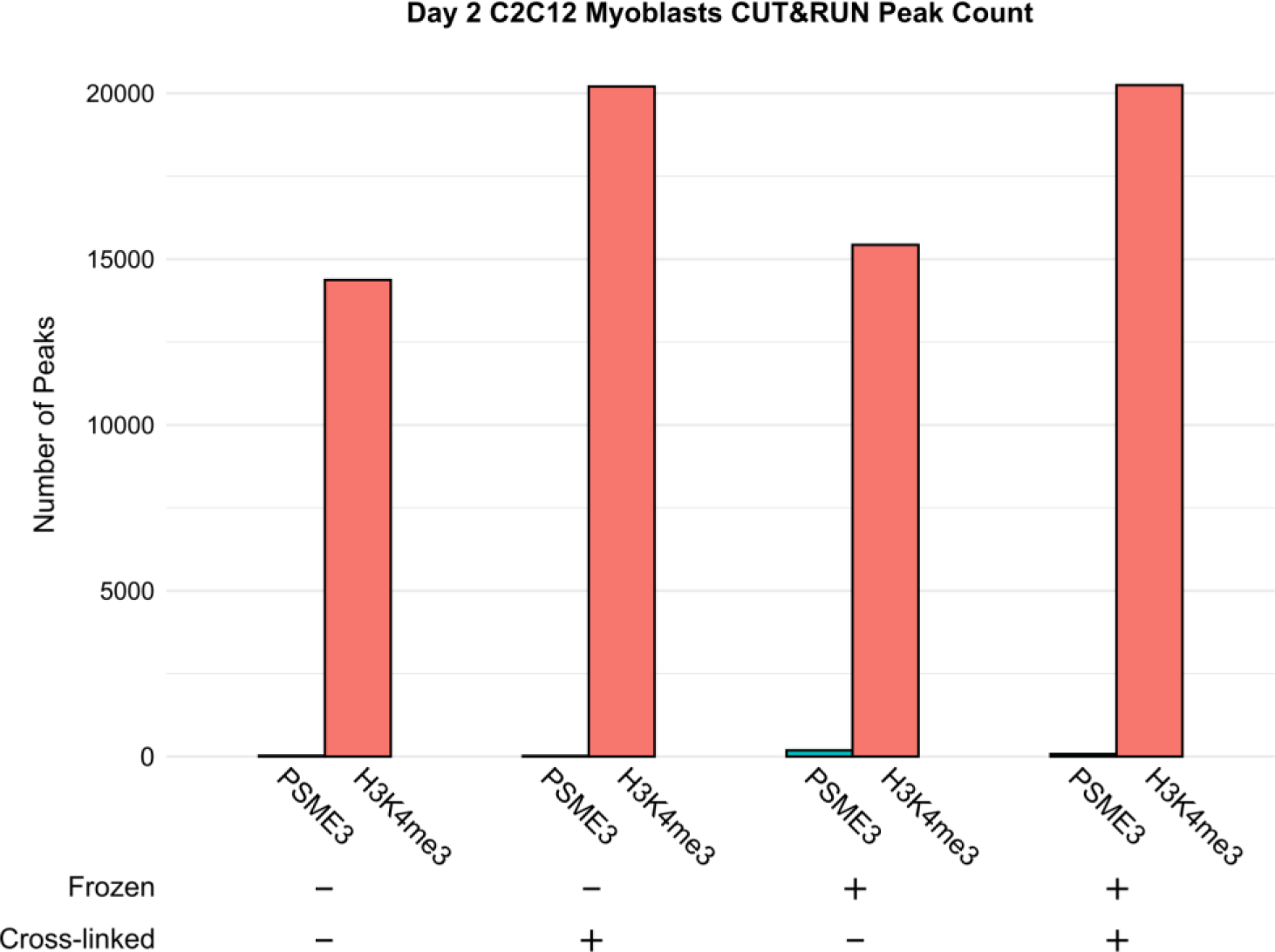
PSME3 is not detected by CUT&RUN in differentiating myoblasts. CUT&RUN was performed using antibodies targeting PSME3 and H3K4me3 in Day 2 C2C12 cells under various preparation conditions. Cells were either assayed fresh after collection, or were first cryopreserved at −80C. Further, cells were either assayed in their native state or after mild cross-linking with formaldehyde.

**Supplementary Figure 2:**
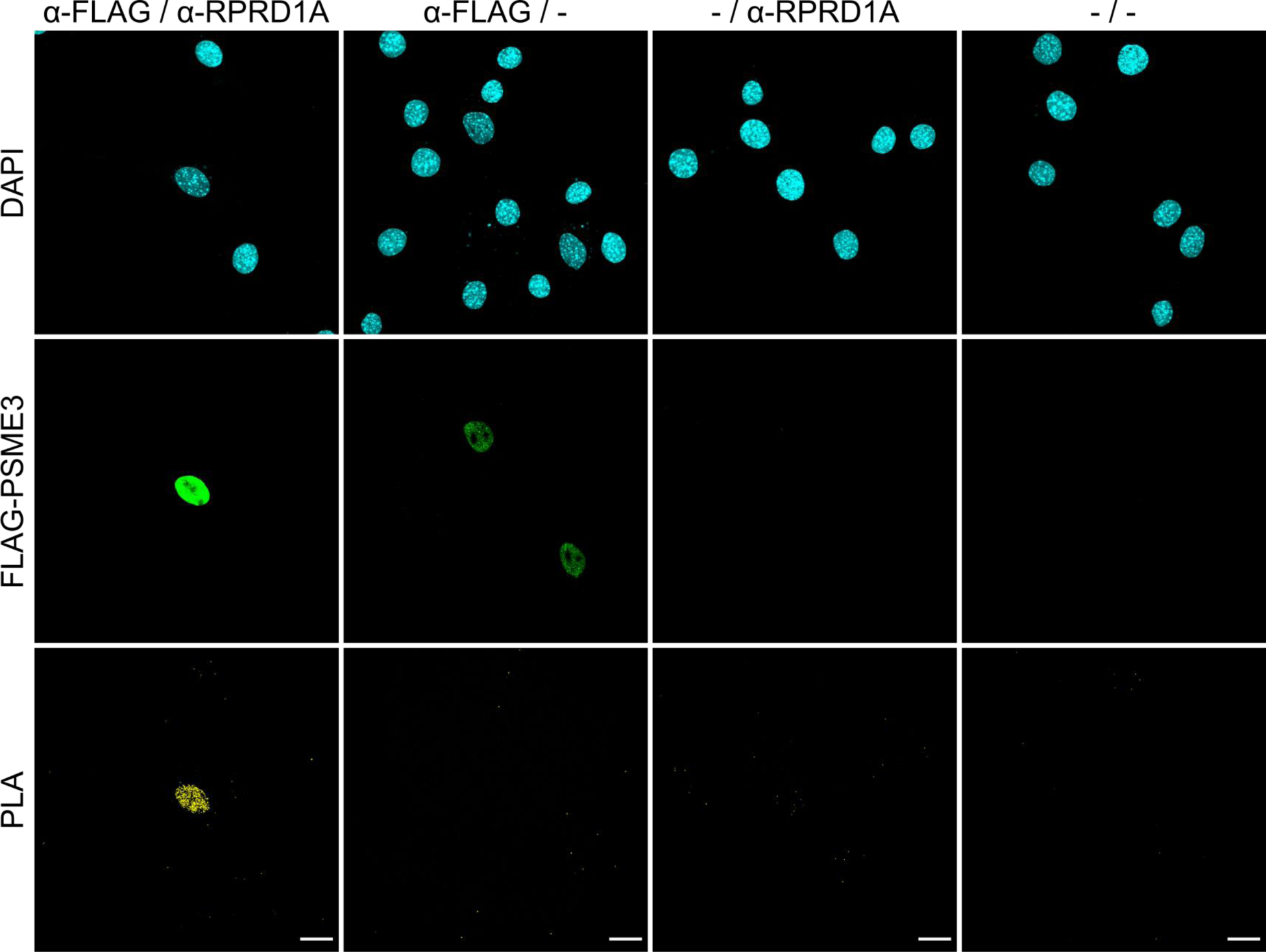
Proximity ligation assay reveals association of PSME3 with RPRD1A. PLA was performed in cells expressing FLAG-PSME3 using antibodies against FLAG and RPRD1A with the respective omission of either or both antibodies as controls. Scale bar is 20 microns in length.

**Supplementary Figure 3:**
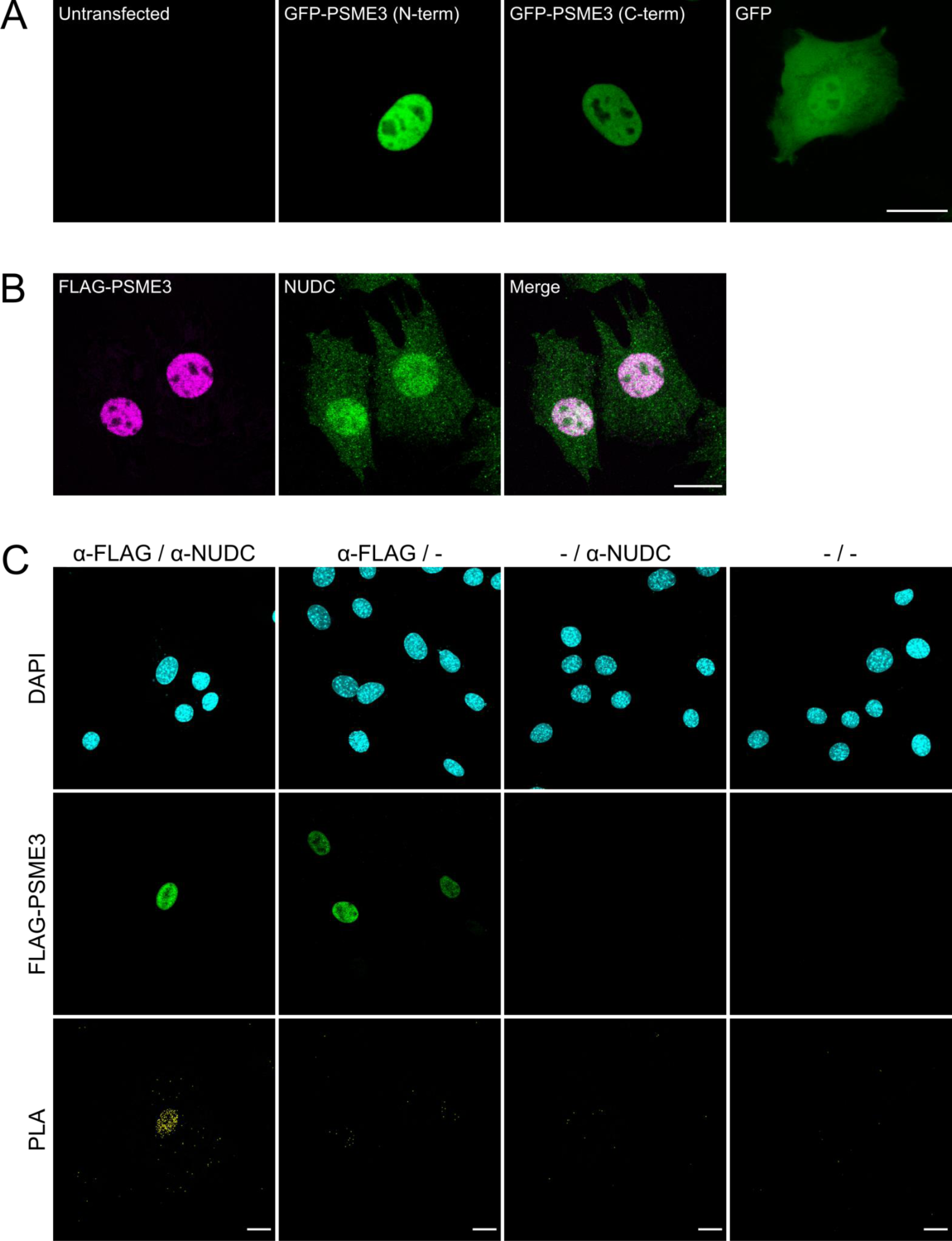
PSME3 and NUDC interact in the nucleus. **A**) Cycling C2C12 cells were transfected with constructs expressing PSME3 tagged with GFP either at its N- or C-terminal, or with a construct expressing free GFP. Cells were imaged live and unstained; scale bar is 20 microns in length. **B**) C2C12 cells expressing FLAG-tagged PSME3 were stained with antibodies against FLAG or endogenous NUDC; scale bar is 20 microns in length. **C**) PLA was performed in cells expressing FLAG-PSME3 using antibodies against FLAG and NUDC with the respective omission of either or both antibodies included as controls. Scale bar is 20 microns in length.

**Supplementary Figure 4:**
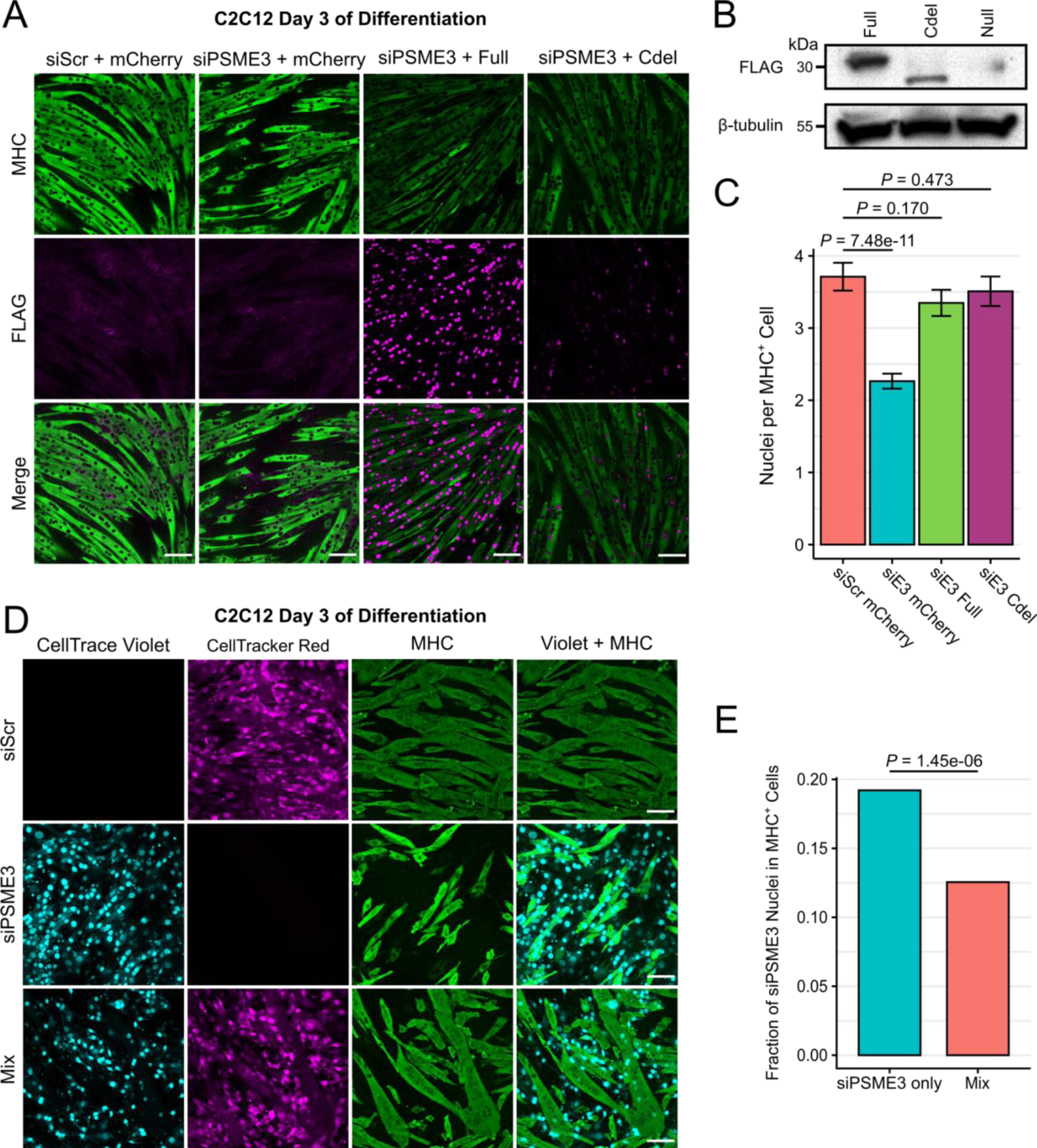
Depletion of PSME3 results in a proteasome-independent, cell-intrinsic differentiation deficit. **A**) Cells were stably transduced with a retroviral construct expressing either mCherry, FLAG-tagged PSME3 (Full), or a truncated FLAG-tagged C-terminal deletion PSME3 mutant (Cdel). Cells were then depleted of endogenous PSME3 by treatment with siRNA and allowed to differentiate for three days before staining. Scale bars are 100 microns in length. **B**) Expression of FLAG constructs in undifferentiated myoblasts. **C**) The number of nuclei contained within each MHC-positive myotube was quantified across conditions and analyzed by the Welch two sample t-test; error bars display the standard error of the mean. **D**) Cells transfected with scrambled (Scr) siRNA or that targeting PSME3 were labeled red or blue, respectively. These cells were induced to differentiate for three days before fixation and staining with an antibody against MHC before imaging. Scale bar is 50 microns. **E**) Quantification of number of siPSME3 (blue) nuclei contained within MHC-positive cells when in the absence or presence of siScr-treated cells; analyzed by two sample t-test of proportions.

